# Electrodetection of small molecules by conformation-mediated signal enhancement

**DOI:** 10.1101/2021.06.17.448904

**Authors:** Krishnan Murugappan, Uthayasuriya Sundaramoorthy, Adam M. Damry, David R. Nisbet, Colin J. Jackson, Antonio Tricoli

**Author notes:** To whom correspondence should be addressed: Colin J. Jackson, Antonio Tricoli. These authors contributed equally to this work (Krishnan Murugappan, Uthayasuriya Sundaramoorthy, Adam M. Damry).

## Abstract

Electrochemical biosensors allow the rapid, selective, and sensitive transduction of critical biological parameters into measurable signals. However, current electrochemical biosensors often fail to selectively and sensitively detect small molecules due to their small size and low molecular complexity. We have developed an electrochemical biosensing platform that harnesses the analyte-dependent conformational change of highly selective solute-binding proteins to amplify the signal generated by analyte binding. Using this platform, we constructed and characterized two biosensors that can sense leucine and glycine, respectively. We show that these biosensors can selectively and sensitively detect their targets over a wide range of concentrations – up to seven orders of magnitude – and that the selectivity of these sensors can be readily altered by switching the bioreceptor’s binding domain. Our work represents a new paradigm for the design of a family of modular electrochemical biosensors, where access to electrode surfaces can be controlled by protein conformational change.

## Introduction

The ability to detect biomolecules rapidly, sensitively, and selectively with portable and small footprint devices is desirable as biomarker concentrations encode a wealth of information regarding metabolic function and health, providing fingerprints for the diagnosis, monitoring, and treatment of diseases.^1-8^ Amongst established biomarker sensing techniques, which include fluorescence, enzyme-linked immunosorbent assays (ELISAs), and surface plasmon resonance, electrochemical approaches offer several desirable advantages such as miniaturization, low cost, and fast response times.^9-11^ Electrochemical sensors are also capable of achieving very high sensitivity, with attomolar limits of detection having been attained by nanostructuring electrode surfaces using metal/metal oxide nanoparticles, carbon nanotubes, graphene, or conducting polymers.^12-14^ Together, these characteristics have led to the widespread adoption of electrochemical sensing platforms in point-of-care (POC) and self-monitoring applications such as blood glucose monitoring for diabetes.^10^

Despite their potential, the performance of electrochemical sensing platforms for sensing small molecule biomarkers (< 1 kDa) is comparatively worse than for macromolecular biomarkers.^11,15,16^ In direct electrochemical sensing platforms, where the analyte is directly oxidized or reduced at the electrode surface, the high redox stability of small molecules requires large overpotentials to activate redox reactions involved in sensing.. Small molecules are also less structurally complex than macromolecules. Together, these factors reduce sensor selectivity.^10,11^ Electrochemical biosensors, which use a biological recognition element coupled to an electrochemical transduction principle, solve this issue of selectivity by relying on the ligand specificity of biomolecules.^10,13,17^ Catalytic biosensors make use of an enzyme or other redox mediators such as metal complexes^11^ to perform a secondary electron transfer reaction with the analyte. However, these biosensors are difficult to generalize (with some exceptions)^18^ and can produce undesirable side-products that can interfere with sensing.^19-21^

In contrast, electrochemical biosensors that use a bioreceptor such as an antibody or aptamer to bind the target of interest to the electrode surface do not require target-specific chemical reactions to function.^17^ Once captured, the target can be detected using either an external redox probe that reports on surface accessibility or an internal probe that reports on bioreceptor morphology.^22^ While these binding-based sensors are effective for sensing large macromolecular targets, the small structural changes induced by small molecule ligand binding are generally not sufficient to allow detection at the required low concentrations.

To overcome the challenges related to current electrochemical small molecule sensing platforms, we have developed a novel approach that can allow selective and sensitive electrochemical detection of small molecules. We demonstrate a new transduction amplification principle that exploits conformational changes in a dynamic bioreceptor protein to enhance the electrochemical response induced by the binding of a small molecule. As proof of this concept, we have engineered solute binding protein (SBP)-based bioreceptors specific to the detection of the amino acids leucine and glycine, two important metabolic biomarkers,^7,8^ and bound them on commercially available screen printed electrodes. The resulting sensing platforms for leucine and glycine are capable of specifically detecting their target ligands over at least five orders of magnitude of concentration and with sensitivities as low as 1 nM and 100 nM, respectively. Given the high specificity of SBPs and their ease of modification toward a target small molecule, our biosensor platform overcomes the issue of poor specificity and the need for complex electrode modifications inherent to previously introduced electrochemical small analyte detection platforms.^11^ In doing so, we have developed the first family of electrochemical sensors that harness a protein conformational change to sensitively detect a small molecule analyte.

## Results

### Design of a binding protein-based electrochemical biosensor

The challenge of designing an electrochemical small molecule biosensor that is modular and generalizable is substantial. To overcome this challenge, we have exploited the large conformation changes that are intrinsic to many small molecule binding proteins, such as the SBP superfamily. Unlike ligand binding to currently widely static bioreceptors, such as antibodies, the small change in relative mass induced by analyte binding to a dynamic bioreceptor can be transduced into a large shape change. This shape change amplifies the change in available electrochemical redox sites on an electrode surface, which is quantified with an external redox probe, providing a tunable and highly sensitive electrochemical platform for sensing small molecules (Fig. 1). Moreover, the shape change and interaction of the binding protein with the electrode surface can be increased or modified by fusing additional rigid domains to a dynamic binding protein core. For example, a protein domain could be fused to the free terminus, which we term the mobile domain, allowing it to be moved through space upon analyte binding to amplify the effect of the conformational change. Alternatively, a protein domain could be inserted between the binding core and the point of immobilization to the electrode surface, which we term a spacer domain, to prevent the solid surface from sterically hindering either analyte binding or the bioreceptor conformational change.

**Figure 1.**
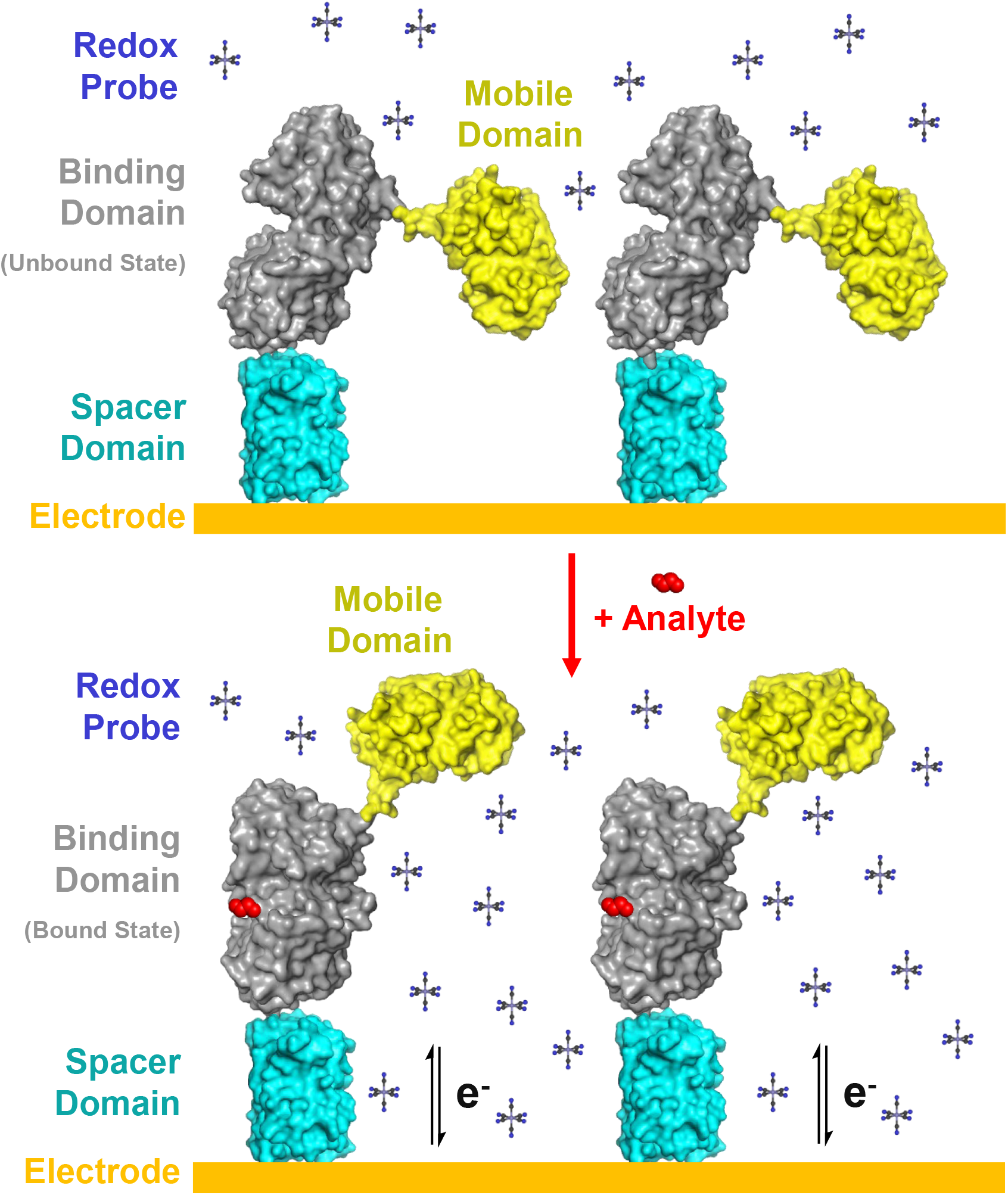
Schematic of the electrochemical sensing mechanism using a dynamic binding protein-based bioreceptor. This bioreceptor consists of three domains; a binding domain (grey) that undergoes a conformational change upon binding of its ligand (red), a mobile domain (yellow) that amplifies the change in protein geometry, and a spacer domain (cyan) that prevents steric hindrance by the electrode surface (gold). Sensing occurs using a redox probe (i.e. ferricyanide, blue) via a conformational change in the binding domain upon analyte binding, altering the morphology of the sensor and its capacity to occlude the electrode surface, thus leading to a change in observed redox probe signal.

To construct this bioreceptor, we reengineered two FRET sensors for use in our electrochemical platform; one based on *E. coli* LivK, a leucine-binding SBP,^23,24^ and the second based on Atu2422 AYW, an engineered glycine-binding SBP previously used to engineer the GlyFS optical biosensor.^25^ These SBPs both undergo a large conformational change upon ligand binding, providing the impetus needed for transduction of the binding event. While fluorescent properties are not necessary for electrochemical sensing, the fluorescent proteins in GlyFS serve as spacer and mobile domains as they are moderately-sized, rigid, inert, and globular proteins. Their use also allows the benchmarking of our constructs’ analyte binding properties using well-established fluorescence-based characterization methods. Therefore, enhanced cyan fluorescent protein (ECFP) was used as a spacer domain and fused directly to the N-terminus of the SBPs, and Venus was used as a mobile domain and fused to the C-terminus using the previously optimized (EAAAK)_3_ rigid linker from GlyFS.^25^ A hexa-histidine tag (His-tag) was fused to the C-terminus of Venus for protein purification using a flexible GGS linker. A tri-cysteine tag (Cys-tag) was also fused to the N-terminus of the ECFP domain using a flexible (GGS)_2_ linker, allowing for covalent attachment of the sensor to the surface of a gold electrode through a sulfhydryl-gold reaction (Supplementary Figs. 1,2). This functionalization strategy allows for the direct and oriented attachment of the sensor to the electrode surface,^26^ which is an important consideration due to the short range of probe-electrode interactions and the anisotropic nature of bioreceptor-electrode interactions.

**Figure 2.**
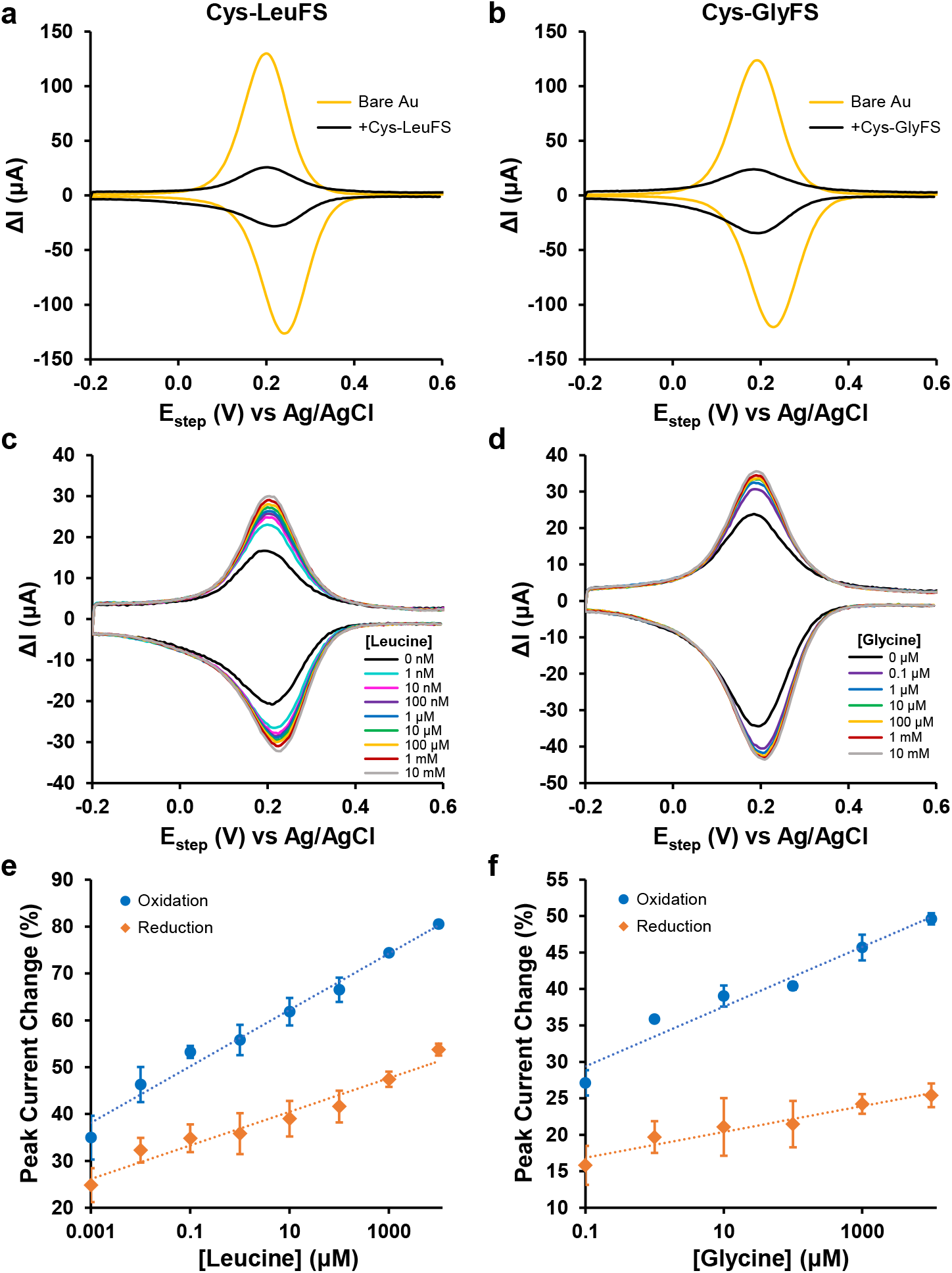
Electrochemical functionalization and sensing responses of Cys-LeuFS and Cys-GlyFS proteins on Au electrodes. Differential pulse voltammograms of Au screen-printed electrodes (Au SPE) functionalized with Cys-LeuFS **(a)** and Cys-GlyFS **(b)** show a decrease in ΔI as compared to bare electrodes. This signal decrease is consistent with a decrease in electron transfer of the ferricyanide (5mM) redox couple due to increased electrode surface occlusion by immobilized protein. Both the Cys-LeuFS **(c)** and Cys-GlyFS **(d)** sensors show an increase in ΔI of 5mM ferricyanide when exposed to different concentrations of glycine and leucine respectively. The increase in redox currents is attributed to the conformational change that arises because of binding of the amino acids to the core of the protein causing the two fluorescent probes to move apart. **(e)** and **(f)** show the calibration plots of both the oxidation and reduction peak currents obtained from **(c)** and **(d)** respectively. All experiments were performed in duplicate, and the average and standard deviation reported for each data point.

To functionalize the electrodes, the two constructs, termed Cys-LeuFS and Cys-GlyFS for the leucine-binding and glycine-binding sensors, respectively, were first heterologously expressed in *Escherichia coli* and purified through a combination of Ni^2+^-NTA affinity chromatography and size-exclusion chromatography (Supplementary Figs. 3,4). Gold screen printed electrodes (Au SPEs) were used because of their low cost and potential for miniaturisation.^27,28^ Following electrode functionalization, we confirmed using scanning electron microscopy that no detectable protein aggregates had deposited on the electrodes (Supplementary Fig. 5). Next, we measured the extent of electrode functionalization with Cys-LeuFS and Cys-GlyFS using differential pulse voltammetry (DPV); a highly sensitive electroanalytical technique.^29,30^ A DPV scan with a bare Au SPE immersed in a ferricyanide solution produced a pair of peaks corresponding to the Fe^2+/3+^ redox couple. In contrast, a reduction in the redox peak currents by 70-80% was observed after attachment of either construct (Fig. 2), which is indicative of a reduction of redox probe diffusion by the bound protein. This decrease in current was stable over several successive scans, which is consistent with the expected covalent binding of the bioreceptor to the electrode.

**Figure 3.**
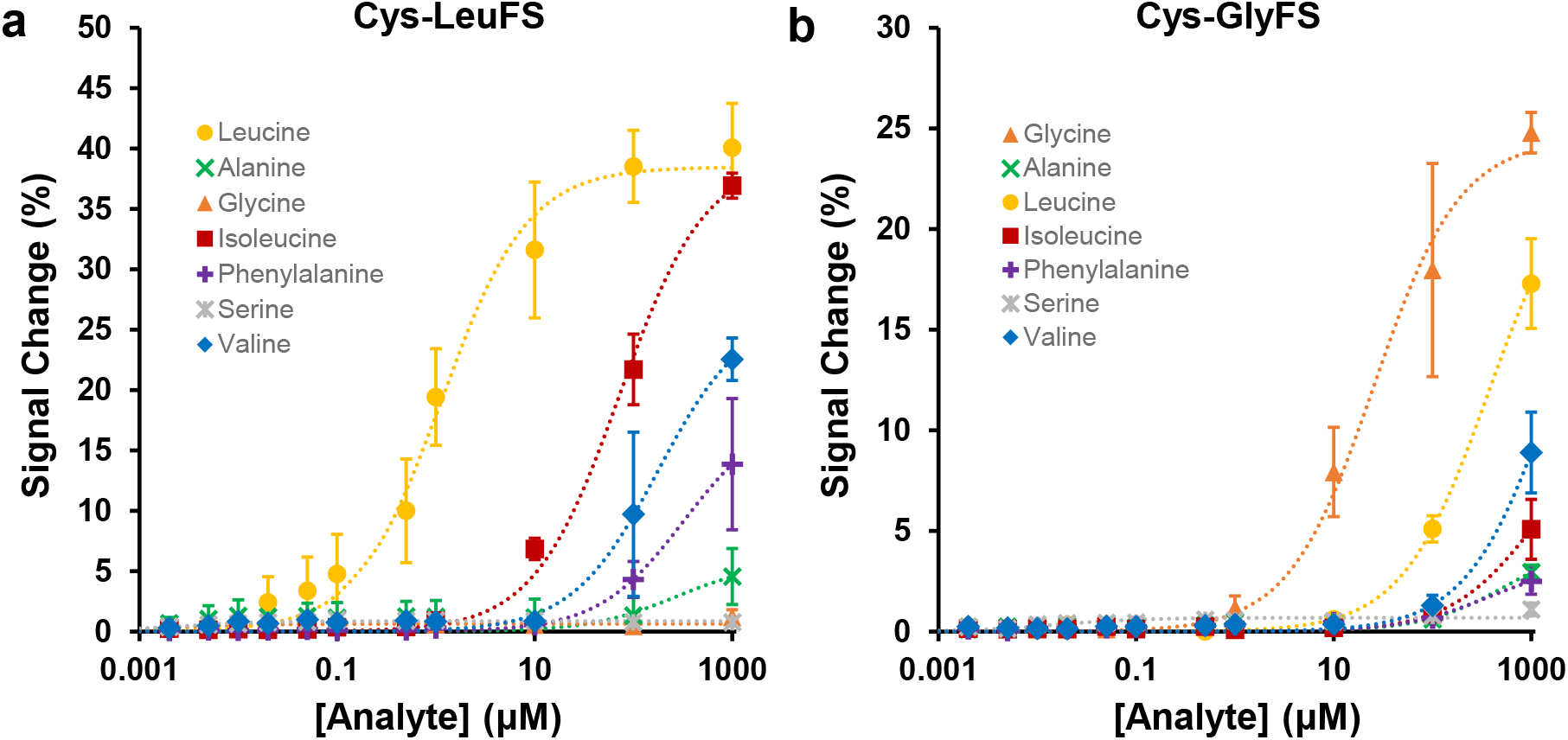
Fluorescence sensing responses of Cys-LeuFS and Cys-GlyFS. Fluorescence dose-response curves of Cys-LeuFS **(a)** and Cys-GlyFS **(b)** to a panel of amino acids including both on- and off-target ligands. 2-term saturation binding models were fit using non-linear regression (dotted lines). Signal change was determined using the ECFP / Venus emission peak ratio (475 nm / 530 nm) relative to signal in absence of any amino acids in solution. All experiments were performed in triplicate, and the average and standard deviation reported for each data point.

**Figure 4.**
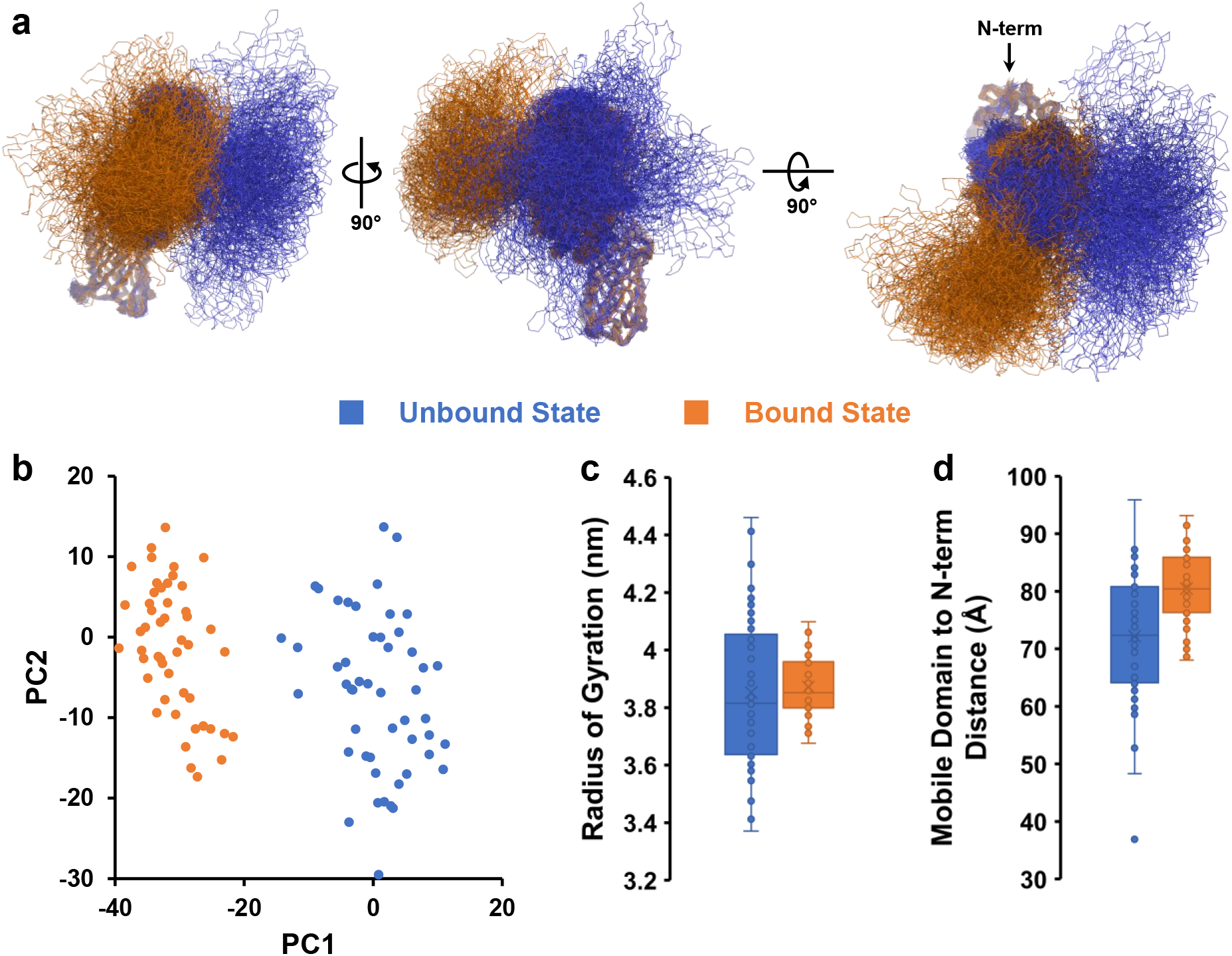
Cys-LeuFS modelling results. **(a)** Conformational ensembles of 50 members each for both the unbound (blue) and ligand-bound (orange) states were generated using molecular dynamics simulations with the MARTINI coarse-grain force field^33^ to equilibrate fusion protein models over 100 ns simulations. Each ensemble member represents the final frame of an independent simulation replicate. The position of the construct N-terminus is indicated in the third orientation to highlight the surface attachment point. **(b)** A principal component analysis of the combined conformational ensemble demonstrates separation of both conformational states along PC1, indicating that both states adopt distinct and separable conformations. **(c)** An analysis of radius of gyration for each ensemble member, shown here as a box plot, shows a broader distribution of radii of gyration for the unbound state in comparison to the ligand-bound state. As radius of gyration is a function of molecular geometry, this is indicative of increased conformational plasticity in the unbound state. **(d)** Box plots of the distance between the mobile domain center of mass to the construct N-terminus (an approximation of distance between the mobile domain and the electrode surface following functionalization) show that the unbound state populates conformations that bring the mobile domain closer to the hypothetical surface location.

**Figure 5.**
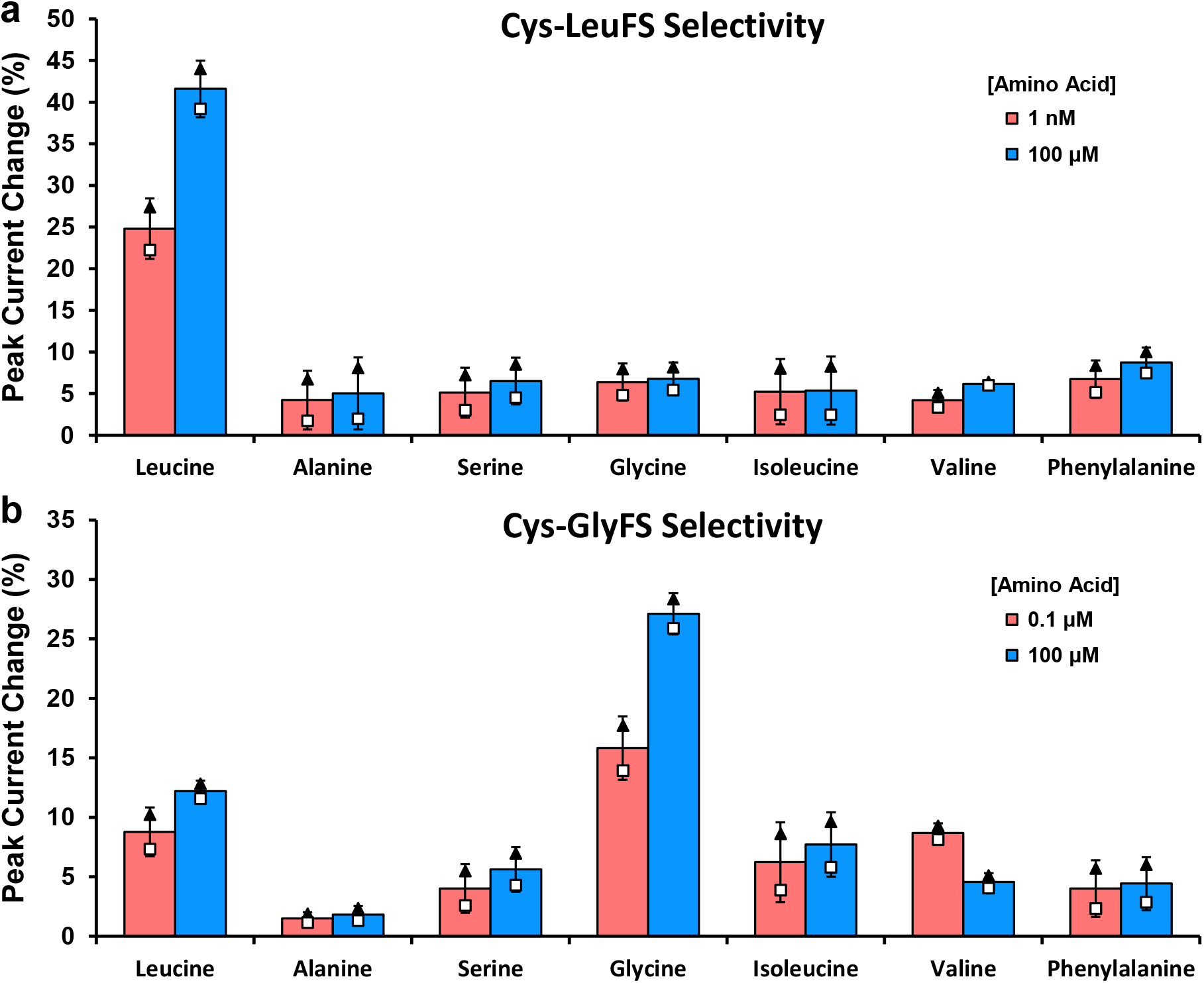
Bar plots depicting selectivity of Cys-LeuFS and Cys-GlyFS against selected amino acids. Selectivity bar plots showing the differential pulse voltammetry response of Cys-LeuFS **(a)** and Cys-GlyFS **(b)** to a panel of 7 free amino acids including both on- and off-target analytes. Signal change was determined at the reduction peak, relative to signal in the absence of any amino acids in solution. All experiments were performed in duplicate, with the bar showing the average of both replicates, error bars representing standard deviation, and results for the two replicates shown separately as black triangles or white squares, respectively.

### Electrochemical sensing using a protein conformational change

The sensitivity of these biosensors was assessed *in vitro* through DPV using ferricyanide solutions spiked with leucine or glycine at concentrations reflecting the solution-state binding affinities of each respective sensor (LivK *K*_D_ (Leu): 8 μM; Atu2422 AYW *K*_D_ (Gly): 20 μM).^24,25^ Exposure of both Cys-LeuFS and Cys-GlyFS to their respective target analytes resulted in an analyte concentration-dependent increase in measured current (Fig. 2c,d), whereas no signal change was observed in a control using unfunctionalized electrodes (Supplementary Fig. 6). The observed change in current is unlikely to have been caused by fouling or by changes in the redox states of leucine and glycine, which are electrochemically inert over the potential ranges used. Thus, the observations of analyte concentration-dependent changes in current when using these sensors are most likely due to the protein conformational changes upon ligand binding increasing electrode surface accessibility. This mechanism is to our knowledge unique amongst current protein-based electrochemical biosensors. A major advantage is the gain-of-signal detection modes induced in comparison to the loss-of-signal modes that are used by many immuno-and DNA-based electrochemical sensors.^9,17^

The dynamic ranges of the sensors were then measured through generation of calibration curves for each sensor. Both sensors exhibited similar relatively log-linear increases in current for both DPV redox peaks over the range of analyte concentrations tested (Fig. 2e,f). The Cys-LeuFS detection range for leucine concentration was shown to range at least seven orders of magnitude, from as low as 1 nM to as high as 10 mM (close to the leucine solubility limit) and with dynamic ranges of at least 50% and 80% for the DPV reduction and oxidation peaks, respectively.

Similarly, Cys-GlyFS was capable of quantitatively sensing glycine concentrations over at least five orders of magnitude, from as low as 100 nM to as high as 10 mM, with dynamic ranges of at least 25% and 50% for the DPV reduction and oxidation peaks, respectively. In contrast, evaluating the performance of these sensors as optical FRET sensors (Fig. 3, Supplementary Fig. 7) showed the expected sigmoidal dose response curves with dynamic ranges of roughly 25-40% and concentration ranges of approximately three orders of magnitude, similar to what has previously been observed for SBP-based FRET biosensors.^24,25^ Both our leucine and glycine sensors therefore display enhanced dynamic ranges and concentration detection ranges when used in electrochemical as opposed to optical platforms. FRET biosensor signaling is dominated by bioreceptor-ligand interactions and therefore follows a simple saturation binding model,^24,25^ whereas electrochemical signaling using these sensors depends on both these bioreceptor-ligand interactions as well as more complex probe diffusion kinetics.

### The molecular basis of the sensing mechanism

To probe the nature of these interactions and understand the molecular mechanism underpinning these sensors’ response, we generated conformational ensembles of Cys-LeuFS in both the leucine-bound and unbound states using an established coarse-grained modelling algorithm (Fig. 4a, Supplementary Figs. 8,9).^31^ The resulting ensembles demonstrate that both states adopt distinct and separable conformational spaces (Fig. 4b). In addition, a solvent exclusion volume analysis^32^ demonstrates that the leucine-bound and unbound states differ in total volume by < 1 % (Supplementary Fig. 10). These results corroborate that a shape effect rather than a volume or size effect is likely responsible for observed signal changes upon analyte binding. Further examining the geometry of these ensemble members shows that while both states exhibit fluctuations of similar magnitudes during equilibration (Supplementary Fig. 11), the protein’s unbound state can adopt a wider range of conformations as evidenced by broader distributions for both radius of gyration (Fig. 4c) and the first four principal component projections (Supplementary Fig. 12). This conformational plasticity could allow protein molecules in the unbound state to adopt conformations that more readily occlude the electrode surface. Indeed, examining the distance between the mobile domain center of mass and the construct’s N-terminus (the location of the surface attachment point), we observe that both the average and minimum distance is considerably reduced for proteins in the unbound state as compared to the bound state (Fig. 4d). This further suggests that in their unbound state, our bioreceptors more strongly inhibit redox probe diffusion to the electrode surface than in their more compact bound state, helping to explain the molecular basis for our sensors’ function.

Given the dynamic nature of this shape-change sensing mechanism, we next verified the reversibility of analyte binding by Cys-GlyFS over repeated measurements by cycling between ferricyanide solutions containing either no glycine or one of two concentrations of glycine (Supplementary Fig. 13). Although signal changes and therefore analyte binding were substantially reversible, a progressive baseline shift was observed over multiple measurements, reaching a plateau after the third return to the glycine-free solution. Despite this baseline shift, the sensor nonetheless remained active and responsive to changes in glycine concentration over all measurements, and the elevated baseline remained below the level of sensor response to even a low glycine concentration. This is attributed to the reversible nature of SBP ligand binding and suggests that the sensor could be used for continuous monitoring applications following additional optimization and/or calibration steps.

### A selective small molecule electrochemical sensor

To demonstrate the excellent and tuneable selectivity of our biosensor platform, we have used amino acids as target analytes as they present a high level of similarity and are known to be very difficult to discriminate between by established approaches.^11^ Thus, the selectivity of our glycine and leucine sensing platforms was assessed against a panel of other common amino acids, including polar, non-polar, small, and bulky amino acids. An initial validation using optical (FRET) sensing showed that Cys-LeuFS was selective for leucine, with some weaker affinity for isoleucine and valine, while Cys-GlyFS was selective for glycine, with some weak affinity for leucine and valine (Fig. 3). We then tested our electrochemical sensing platform’s ability to selectively sense these amino acids using the same panel of amino acids at both a low concentration (1 nM for Cys-LeuFS and 100 nM for Cys-GlyFS) and a high concentration (100 μM for both sensors) (Fig. 5, Supplementary Figs. 14,15). The Cys-LeuFS-functionalized sensors demonstrated high selectivity for leucine, with the off-target signal observed at high concentrations of isoleucine or valine in optical sensing being largely absent in electrochemical sensing.

The Cys-GlyFS-functionalized sensors displayed the highest response for glycine, with some affinity for leucine and valine. While the trend in terms of selectivity is consistent with the FRET data, it is notable that the sensitivity appears to be greater when the proteins are used as part of our electrochemical sensing platform; a > 20% current change for Cys-LeuFS in the presence of 1 nM Leucine, with selective detection over seven orders of magnitude (1 nM -10 mM) is remarkable. Most importantly, the ligand binding domain of these sensors is the only domain that differs between both platforms. We can thus conclude that the different selectivity observed for these sensors (Cys-LeuFS *vs*. Cys-GlyFS) indicates that the platform’s selectivity is dictated by the ligand binding domain and that swapping this domain for another, in a modular fashion, with a different ligand selectivity is sufficient to alter the selectivity of the biosensor.

## Discussion

In this work, we developed a novel protein-based electrochemical biosensor platform, exemplified by two amino acid biosensors: Cys-LeuFS and Cys-GlyFS, which are, to our knowledge, the first non-catalytic electrochemical biosensors that use a protein conformational change as transduction mechanism to detect a small molecule. This was accomplished by reengineering two SBP-based optical biosensors, resulting in a marked increase in the sensors’ dynamic ranges and sensitivity upon translation to an electrochemical detection platform. These sensors’ properties can solve several major issues in electrochemical sensing of small molecules, where large overpotentials and low selectivity are commonplace, rendering many existing electrochemical sensors unsuitable to clinical applications.^11^ In addition to these direct advantages in sensing, the modular nature of these novel sensors, where the core bioreceptor dictates selectivity, makes them appealing for future engineering and diversification efforts. Notably, this work lays out the foundation for readily creating a family of other SBP-based electrochemical biosensors^34-37^. This process is facilitated by the less strict geometries required in electrochemical sensing relative to the optimization of FRET efficiency changes in optical sensing, the latter of which often requires extensive linker optimization.^25^ The ability to directly translate optical FRET sensors to electrochemical sensors also benefits efforts to further engineer sensor selectivity, as optical sensors are particularly amenable to protein engineering efforts due to the availability of high-throughput optical screening methods.^38,39^ Therefore, our available toolbox of electrochemical biosensors could readily be expanded through engineering optical SBP-based sensors with altered ligand selectivity or affinity, then translating these into highly sensitive electrochemical platforms.

Beyond modifications to our biosensors’ selectivity, considerable room for future optimization of these sensors’ response remains. Optimizing the size, shape, and net charge of their mobile and static domains, consequently changing the electrochemical morphology of the sensor, would further enhance the signal change observed upon analyte binding. In addition, the current iteration of these sensors requires the use of an external redox probe to generate the currents measured. An internal redox probe could instead be attached to the mobile region of the sensor, creating a fully self-contained electrochemical biosensing platform. Despite these avenues for future improvement, the ability of these protein-based bioreceptors to selectively capture and detect small molecule targets is already a significant advancement compared to current methods that rely on using electrocatalysts or redox mediators. Both of these existing approaches can lack selectivity, especially towards structurally simple molecules such as glycine.^11^ Cys-LeuFS and Cys-GlyFS are instead capable of discriminating their respective native ligands leucine and glycine from other chemically-similar chain amino acids, highlighting the exquisite specificity that can be attained using binding proteins as bioreceptors. Overall, we have shown how highly sensitive and specific electrochemical biosensors can be developed by integrating dynamic binding protein-based bioreceptors with highly sensitive electro-analytical techniques. This novel platform opens avenues for the detection of small molecules such as leucine and glycine, two medically relevant amino acids that are involved in key biological functions yet can be difficult to sense using small footprint sensing platforms. Future optimization of our biosensors’ protein elements and the ready miniaturization of the platform enabled using SPEs bears potential for a large range of applications, including their use as miniaturized sensors for portable and point of care medical diagnostic, and environmental monitoring.

## Materials and Methods

### Gene constructs and cloning

Cys-LeuFS and Cys-GlyFS constructs were designed by sequential fusion of three protein domains. The N-terminal domain consists of ECFP with an N-terminal Cys-tag and a 9 amino acid truncation at its C-terminus to remove the flexible ECFP C-terminal tail. This ECFP construct was fused to a leucine-binding (*E. coli* LivK) or glycine-binding (Atu2422 AYW) solute binding protein from which the signal peptide was truncated, which was in turn fused to a C-terminal Venus domain using the rigid linker (EAAAK)_3_. Codon-optimized genes for Cys-LeuFS and Cys-GlyFS cloned into the pET-29b(+) and pET-28a(+) vectors respectively were obtained from Twist Biosciences and transformed via electroporation into *E. coli* BL21(DE3).

### Protein expression and purification

Proteins were expressed using an autoinduction medium (yeast extract, 5 g/L; tryptone, 20 g/L; NaCl, 5 g/L; KH_2_PO_4_, 3 g/L; Na_2_HPO_4,_ 6 g/L with 10 ml/L 60 % (v/v) glycerol, 5 ml/L 10 % (w/v) glucose, 25 ml/L 8 % (w/v) lactose) supplemented with 50 mg kanamycin or 100 mg ampicillin for Cys-LeuFS or Cys-GlyFS expression, respectively. Cultures were grown at 18 °C with shaking for 72-96 hours with periodic monitoring of ECFP and Venus fluorescence to gauge protein expression. Following growth, cells were harvested by centrifugation and stored at -20 °C. For purification, the frozen pellet was suspended in buffer A (50 mM sodium phosphate, 300 mM NaCl, 20 mM imidazole, pH 7.4), lysed by sonication, re-centrifuged at high speed (22400 xg for 60 min at 4 ºC) and the cleared supernatant was collected. This was loaded onto a 5 mL Ni-NTA/His-trap column pre-equilibrated in buffer A, washed with 10 column volumes of buffer A, followed by 5 column volumes of 10 % buffer B, and eluted with 100 % buffer B (50 mM sodium phosphate, 300 mM NaCl, 250 mM imidazole, pH 7.4), and the eluted sensor protein was dialyzed against 3 exchanges of 4 L of buffer C (20 mM sodium phosphate, 200 mM NaCl, pH 7.4) at 4ºC. The dialyzed protein was further purified using a GE Healthcare HiLoad 26/600 Superdex 200 pg SEC column using buffer C. Protein purity was confirmed by SDS-PAGE, and protein concentrations were measured spectrophotometrically using predicted molar absorption coefficients.

### Fluorescence assays

Fluorescence titrations were performed using a Cary Eclipse fluorimeter (Varian) using a 1 cm quartz narrow volume fluorescence cuvette (Hellma Analytics). Protein samples containing 2 uM Cys-LeuFS or Cys-GlyFS and 0.001 μM to 1000 μM free amino acids (leucine, glycine, alanine, valine, isoleucine, serine, and phenylalanine) in buffer C (20 mM sodium phosphate, 200 mM NaCl, pH 7.4) underwent excitation at 433 nm and emission scans from 450 nm to 560 nm were obtained in triplicate with a step size of 1 nm. ECFP/Venus fluorescence ratios were determined using fluorescence intensities at 475 nm (ECFP emission peak) and 530 nm (Venus emission peak). *K*_*D*_ values were determined by fitting curves through non-linear regression using a saturation binding model:

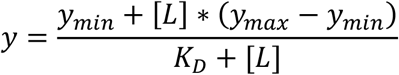

where *[L]* represents concentration of ligand in solution, and *y* represents fluorescent signal.

### Electrochemical methods

All electrochemical experiments were performed using a VMP3 potentiostat (BioLogic) interfaced to a PC with EC-Lab software. Au SPEs (Au AT, Dropsens) were first cleaned in 0.1 M H_2_SO_4_ (Sigma-Aldrich) by cycling between 0 and 1.4 V for 10 times at a scan rate of 1 V/s. Following this they were dipped in a solution of 5 mM K_3_Fe(CN)_6_ (Sigma-Aldrich) in phosphate buffered saline (20 mM sodium phosphate, 200 mM NaCl, pH 7.4) and scanned between the potential range -0.2 to 0.6 V vs Ag/AgCl using differential pulse voltammetry (DPV) with an amplitude of 0.05 V, modulation time of 0.05 s and interval time of 0.1 s. Once stable redox peaks characteristic of the Fe^2+/3+^ couple were seen, the SPEs were removed, washed with DI and then dried before drop casting 100 μl of Cys-LeuFS/Cys-GlyFS for 3 hours. This functionalized surface was dipped back into 5 mM K_3_Fe(CN)_6_/0.1 M PBS to record following DPVs. The electrodes were then exposed to different concentrations of amino acids (leucine, glycine, alanine, valine, isoleucine, serine, or phenylalanine, from Sigma-Aldrich) with DI washing steps before and after every exposure.

### Scanning Electron Microscopy

The morphology of the Au screen printed electrodes was investigated using a field-emission scanning electron microscope Zeiss Ultra Plus (operating at 3 kV) without coating.

### Construct modelling

Starting structures were generated in PyMOL^40^ by fusing unbound and ligand-bound LivK (PDB ID: 1USG and 1USK, respectively)^41^ directly to ECFP (PDB ID: 5OX8)^42^ and to Venus (PDB ID: 1MYW)^43^ using a fully-extended (EAAAK)_3_ linker. Coarse grain simulations were prepared by solvating starting structures in a dodecahedral solvent box with a minimum distance of 10 Å from any protein atom to the box wall followed by coarse graining using the Martinize script. Coarse grain simulations were run in GROMACS^44^ using the MARTINI forcefield.^33^ Structures were first minimized using steepest-descent energy minimization, followed by a 100 ps equilibration in the NVT ensemble and a 100 ps equilibration in the NPT ensemble. Structure equilibration including linker collapse was carried out over 50 replicate trajectories of 100 ns for each state, with a 2.5 fs timestep and an elastic network model used to maintain domain geometry with restraints on linker beads removed to allow free equilibration of the linker. Temperature coupling used a V-rescale thermostat and pressure coupling used a Parinello-Rahman barostat. Following equilibration, coarse grain structures were converted back to fine grain models using the Backwards tool.^45^ Geometry analysis was carried out in VMD,^46^ solvent exclusion volume was calculated using ProteinVolume 1.3,^32^ and PCA analysis was carried out using PRODY.^47^

## Data Availability

The main data supporting the findings of this study are available within the paper and its supplementary information. Any other relevant data are available from the corresponding authors upon reasonable request. Source data are provided with this paper.

## Acknowledgements

This research was funded by and has been delivered in partnership with Our Health in Our Hands (OHIOH), a strategic initiative of the Australian National University, which aims to transform healthcare by developing new personalised health technologies and solutions in collaboration with patients, clinicians, and health care providers. A.T. gratefully acknowledges the support of the Australian Research Council for a Future Fellowship (FT200100939) and a Discovery grant DP190101864, and from the North Atlantic Treaty Organization Science for Peace and Security Programme project AMOXES (#G5634). D.R.N was supported by a NHMRC Research Leadership Fellowship (GNT1135657). A.M.D. acknowledges funding from the Human Frontier Science Program (LT-000366/2020-C). Funding by the Australian Research Council Centre of Excellence for Innovations in Peptide and Protein Science and the Centre of Excellence in Synthetic Biology is gratefully acknowledged. This project was undertaken with the assistance of resources and services from the National Computational Infrastructure (NCI), which is supported by the Australian Government.

## Author Contributions

A.M.D., A.T., C.J.J., D.N., and K.M. conceptualized the electrochemical platform. A.M.D., C.J.J., and U.S. designed and optimized the platform’s protein elements. A.T. and K.M. designed and optimized the platform’s electrochemical elements. U.S. expressed protein and performed and analyzed fluorescence characterization experiments. K.M. performed scanning electron microscopy experiments, performed electrochemical sensing experiments, and analyzed electrochemical sensing data. A.M.D. generated and analyzed bioreceptor computational models. A.M.D., A.T., C.J.J., and D.N. obtained funding for the project. A.M.D., A.T., C.J.J., and K.M. wrote the manuscript. All authors reviewed the manuscript.

