## Supplementary Information for "Electrodetection of small molecules by conformation-mediated signal enhancement"

* To whom correspondence should be addressed:

1 10 20 30 40 50 60 70
MCCC**GGSGGS**MVSKGEELFTGVVPILVELDGDVNGHKFSVSGEGEGDATYGKLTLKFICTTGKLPVPWPT
71 80 90 100 110 120 130 140
LVTTLTWGVQCFSRYPDHMKQHDFFKSAMPEGYVQERTIFFKDDGNYKTRAEVKFEGDTLVNRIELKGID
141 150 160 170 180 190 200 210
FKEDGNILGHKLEYNYISHNVYITADKQKNGIKANFKIRHNIEDGSVQLADHYQQNTPIGDGPVLLPDNH
211 220 230 240 250 260 270 280
YLSTQSALSKDPNEKRDHMVLLEFVTAAGIIKVAVVGAMSGPIAQWGDMEFNGARQAIKDINAKGGIKGD
281 290 300 310 320 330 340 350
KLVGVEYDDACDPKQAVAVANKIVNDGIKYVIGHLCSSSTQPASDIYEDEGILMISPGATNPELTQRGYQ
351 360 370 380 390 400 410 420
HIMRTAGLDSSQGPTAAKYILETVKPQRIAIIHDKQQYGEGLARSVQDGLKAANANVVFFDGITAGEKDF
421 430 440 450 460 470 480 490
SALIARLKKENIDFVYYGGYYPEMGQMLRQARSVGLKTQFMGPEGVGNASLSNIAGDAAEGMLVTMPKRY
491 500 510 520 530 540 550 560
DQDPANQGIVDALKADKKDPSGPYVWITYAAVQSLATALERTGSDEPLALVKDLKANGANTVIGPLNWDE
561 570 580 590 600 610 620 630
KGDLKGFDFGVFQWHADGSSTAAK**EAAAKEAAAKEAAAK**VSKGEELFTGVVPILVELDGDVNGHKFSVSG
631 640 650 660 670 680 690 700
EGEGDATYGKLTLKLICTTGKLPVPWPTLVTTLGYGLQCFARYPDHMKQHDFFKSAMPEGYVQERTIFFK
701 710 720 730 740 750 760 770
DDGNYKTRAEVKFEGDTLVNRIELKGIDFKEDGNILGHKLEYNYNSHNVYITADKQKNGIKANFKIRHNI
771 780 790 800 810 820 830 840
EDGGVQLADHYQQNTPIGDGPVLLPDNHYLSYQSALSKDPNEKRDHMVLLEFVTAAGITLGMDELYK**GGS**
841 846
HHHHHH

Cys-Tag ECFP Leucine Binding Protein Venus His-Tag

**Linkers (Flexible + Rigid)**

**Figure S1.** Cys-LeuFS amino acid sequence with the ECFP, Leucine Binding Protein, and Venus domains highlighted in cyan, grey, and yellow, respectively. The N-terminal Cys-tag and C-terminal His-tag are highlighted in magenta and green, respectively. Linkers are shown in bold, with the rigid linker between the Leucine Binding Protein and Venus domains additionally underlined.

Cys-GlyFS

1 10 20 30 40 50 60 70

MCCC**GGSGGS**MVSKGEELFTGVVPILVELDGDVNGHKFSVSGEGEGDATYGKLTLKFICTTGKLPVPWPT

71 80 90 100 110 120 130 140

LVTTLTWGVQCFSRYPDHMKQHDFFKSAMPEGYVQERTIFFKDDGNYKTRAEVKFEGDTLVNRIELKGID

141 150 160 170 180 190 200 210

FKEDGNILGHKLEYNYISHNVYITADKQKNGIKANFKIRHNIEDGSVQLADHYQQNTPIGDGPVLLPDNH

211 220 230 240 250 260 270 280

YLSTQSALSKDPNEKRDHMVLLEFVTAAGIDVVIAVGAPLTGPNAAFGAQIQKGAEQAAKDINAAGGING

281 290 300 310 320 330 340 350

EQIKIVLGDDVSDPKQGISVANKFVADGVKFVVGHANSGVSIPASEVYAENGILEITPYATNPVFTERGL

351 360 370 380 390 400 410 420

WNTFRTCGRDDQQGGIAGKYLADHFKDAKVAIIHDKTPYGQGLADETKKAANAAGVTEVMYEGVNVGDKD

421 430 440 450 460 470 480 490

FSALISKMKEAGVSIIYWGGWHTEAGLIIRQAADQGLKAKLVSGDGIVSNELASIAGDAVEGTLNTFGPD

491 500 510 520 530 540 550 560

PTLRPENKELVEKFKAAGFNPEAYTLYSYAAMQAIAGAAKAAGSVEPEKVAEALKKGSFPTALGEISFDE

561 570 580 590 600 610 620 630

KGDPKLPGYVMYEWKKGPDGKFTYIQQ**EAAAKEAAAKEAAAK**VSKGEELFTGVVPILVELDGDVNGHKFS

631 640 650 660 670 680 690 700

VSGEGEGDATYGKLTLKLICTTGKLPVPWPTLVTTLGYGLQCFARYPDHMKQHDFFKSAMPEGYVQERTI

701 710 720 730 740 750 760 770

FFKDDGNYKTRAEVKFEGDTLVNRIELKGIDFKEDGNILGHKLEYNYNSHNVYITADKQKNGIKANFKIR

771 780 790 800 810 820 830 840

HNIEDGGVQLADHYQQNTPIGDGPVLLPDNHYLSYQSALSKDPNEKRDHMVLLEFVTAAGITLGMDELYK

841 849

**GGS**HHHHHH

Cys-Tag ECFP Glycine Binding Protein Venus His-Tag

**Linkers (Flexible + Rigid)**

**Figure S2.** Cys-GlyFS amino acid sequence with the ECFP, Glycine Binding Protein, and Venus domains highlighted in cyan, grey, and yellow, respectively. The N-terminal Cys-tag and C-terminal His-tag are highlighted in magenta and green, respectively. Linkers are shown in bold, with the rigid linker between the Glycine Binding Protein and Venus domains additionally underlined.

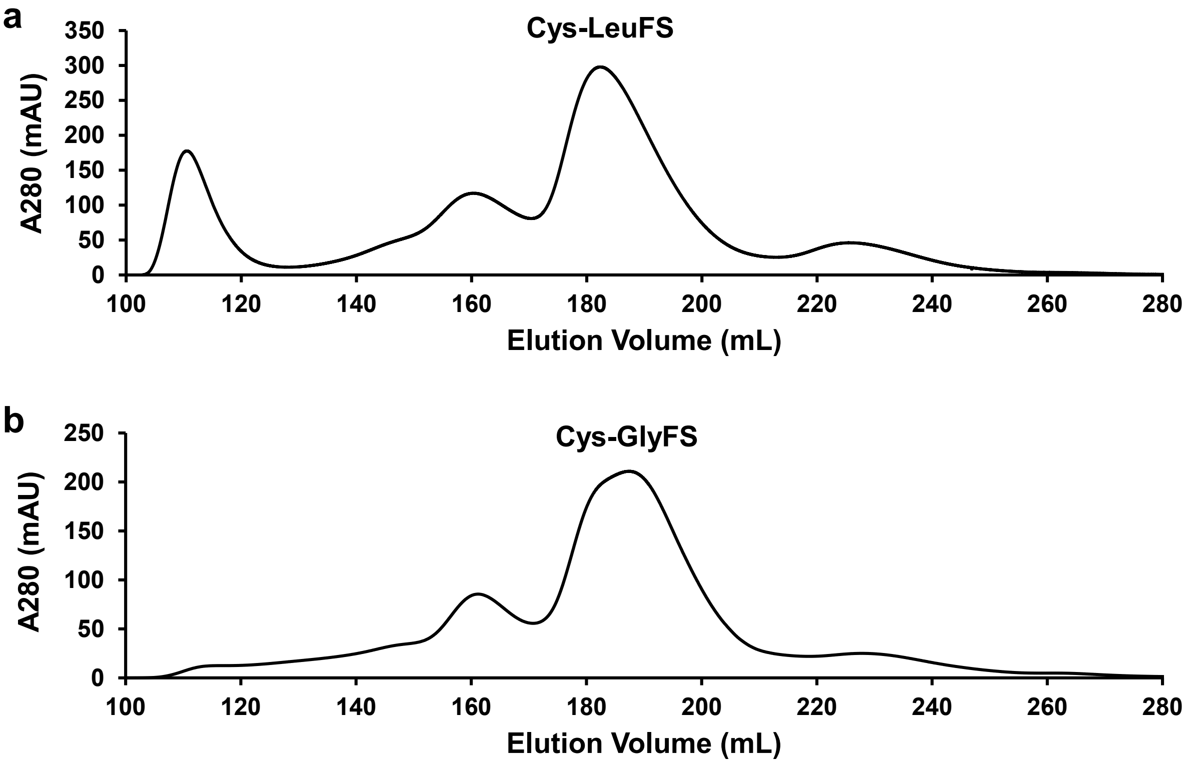


**Figure S3.** Size-exclusion chromatography profiles of Ni-NTA purified Cys-LeuFS (**a**) and Cys-GlyFS (**b**) using an AKTA HiLoad 26/600 Superdex 200 column (column volume = 108 mL). Both proteins elute primarily as monomers at an elution volume of approximately 180 mL, with an additional dimer fraction eluting at an elution volume of approximately 160 mL.

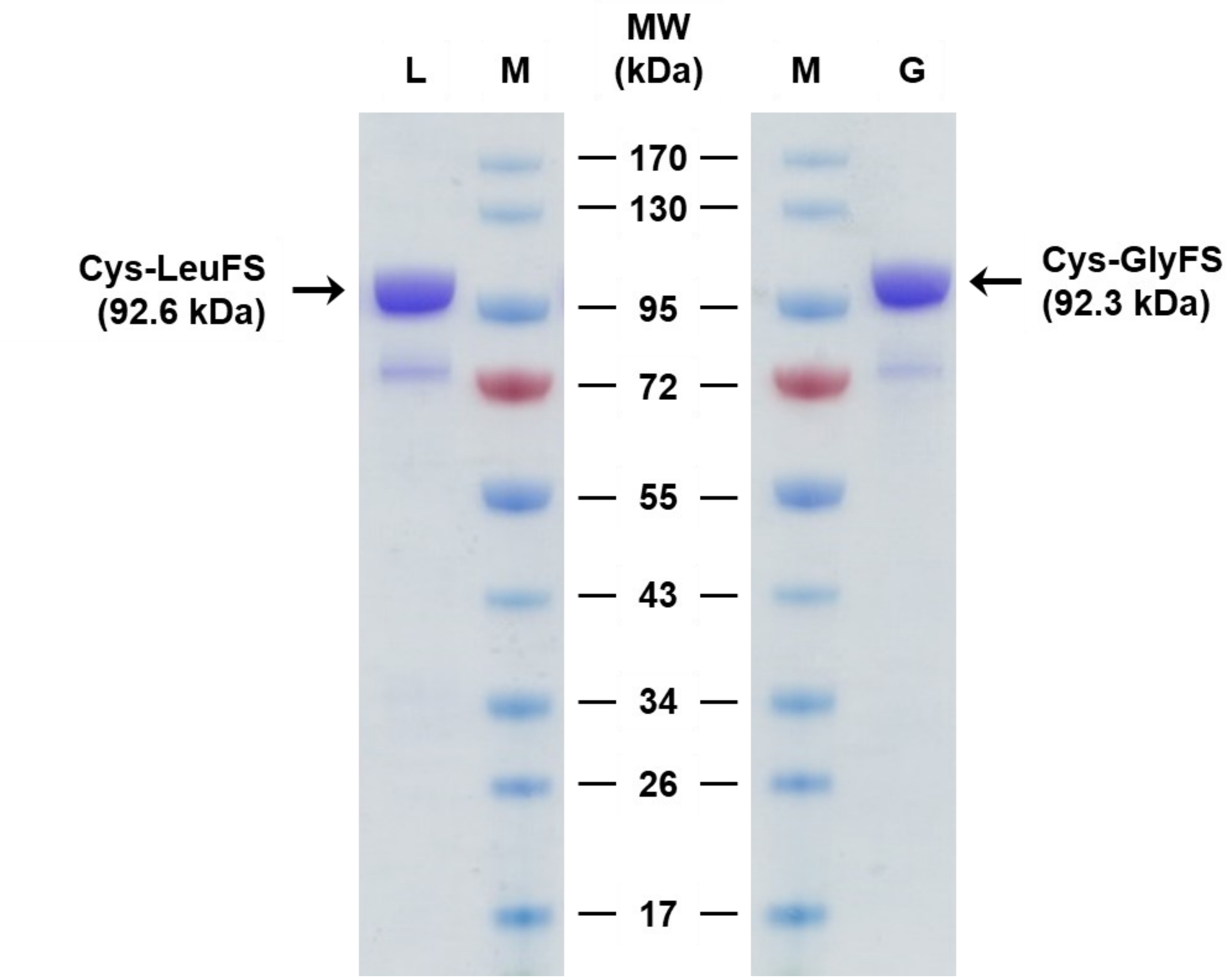


**Figure S4.** SDS-PAGE of size exclusion chromatography-purified Cys-LeuFS and Cys-GlyFS. 1 μg of protein was loaded in a Genscript ExpressPlus 4-20% gradient gel under reducing conditions. M: molecular weight marker (Fisher PageRuler Prestained Protein Ladder). L: Cys-LeuFS. G: Cys-GlyFS. The gel was stained with Coomassie Blue R-250 and destained with methanol:acetic acid:water (4 : 1 : 5, v/v).

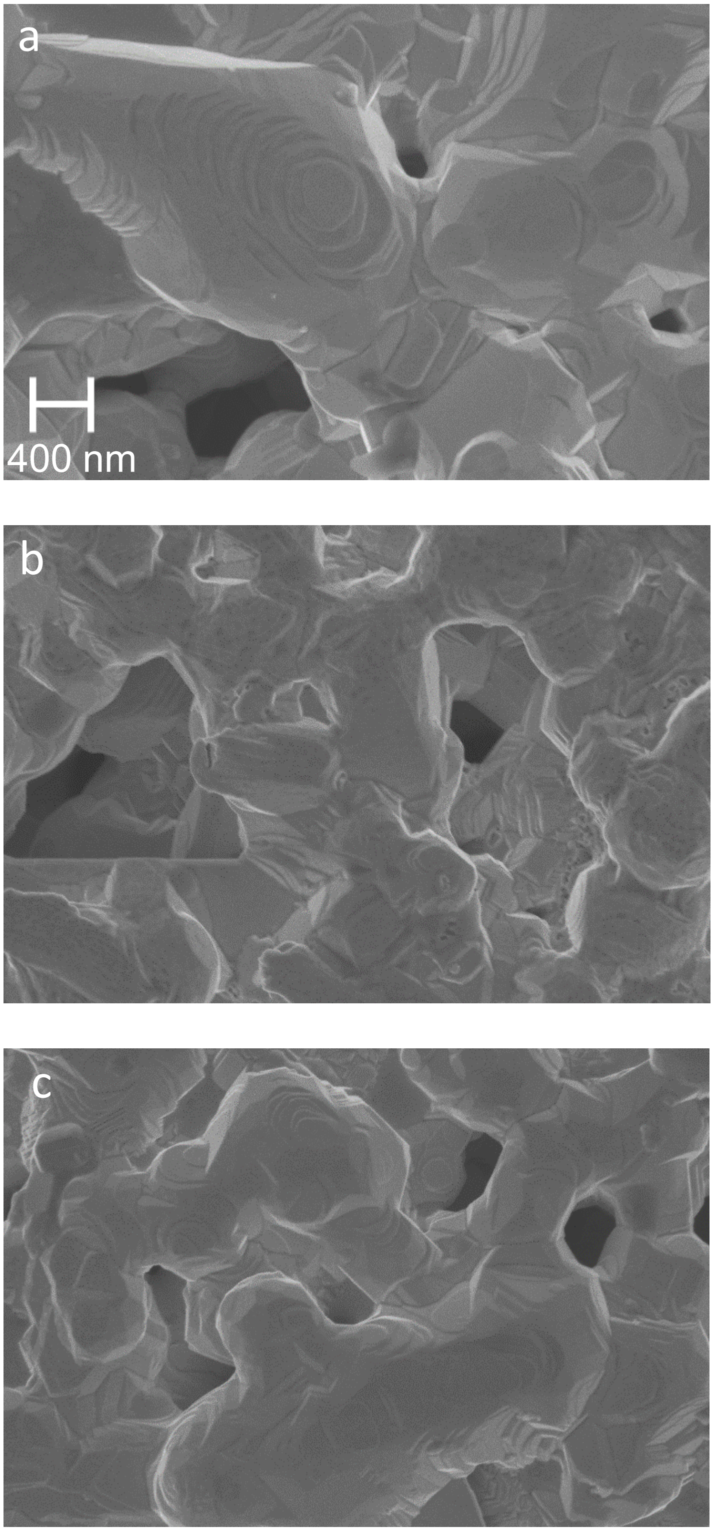


**Figure S5.** Scanning electron micrographs of **(a)** a bare Au SPE, **(b)** a Cys-GlyFS functionalized Au SPE, and **(c)** a Cys-LeuFS functionalized Au SPE.

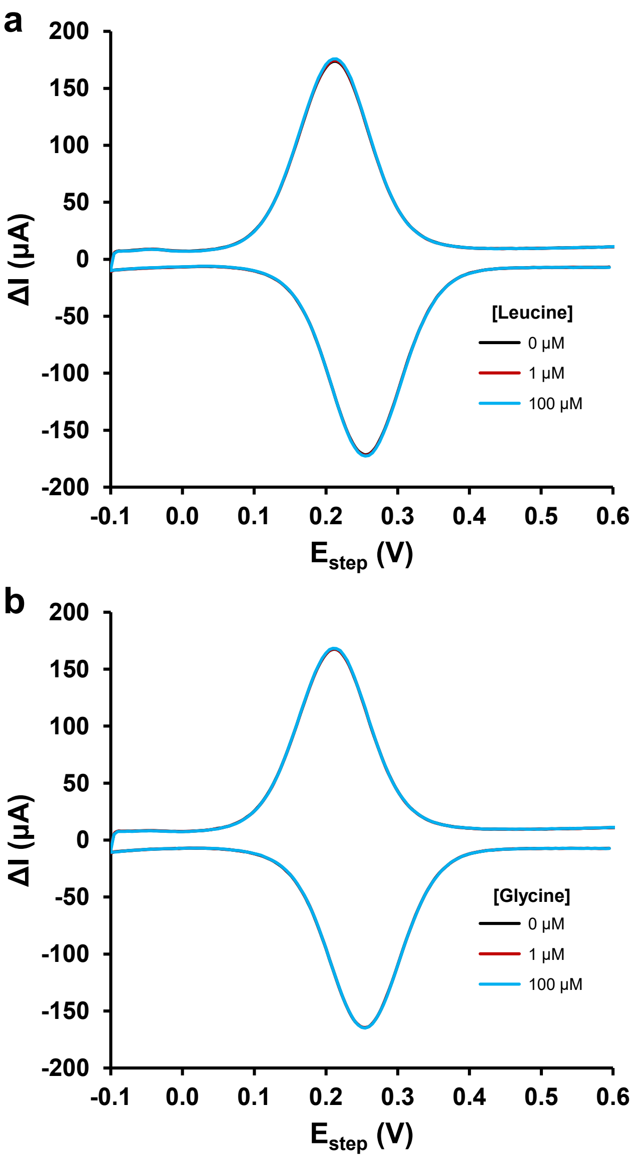


**Figure S6.** Differential pulse voltammograms of bare Au screen-printed electrodes (Au SPE) electrodes in 5 mM potassium ferricyanide show no changes in electrochemical response when leucine **(a)** or glycine **(b)** are added to the solution in the absence of Cys-GlyFS and Cys-LeuFS.

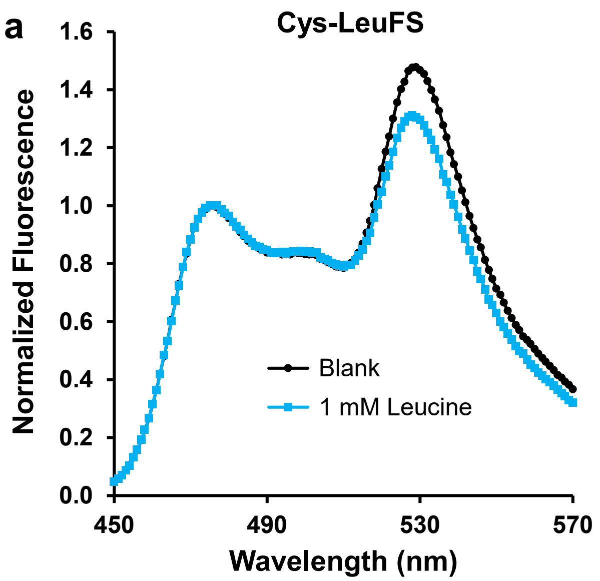


**Figure S7.** Characteristic fluorescence spectra of Cys-LeuFS (**a**) and Cys-GlyFS (**b**) with excitation at 433 nm demonstrate the double-peak arising from ECFP and Venus fluorescence. Substrate binding by the Leucine Binding Protein or Glycine Binding Protein domains reduces FRET efficiency between the ECFP donor and Venus acceptor fluorophores, altering the ratio between the two fluorescence peaks.

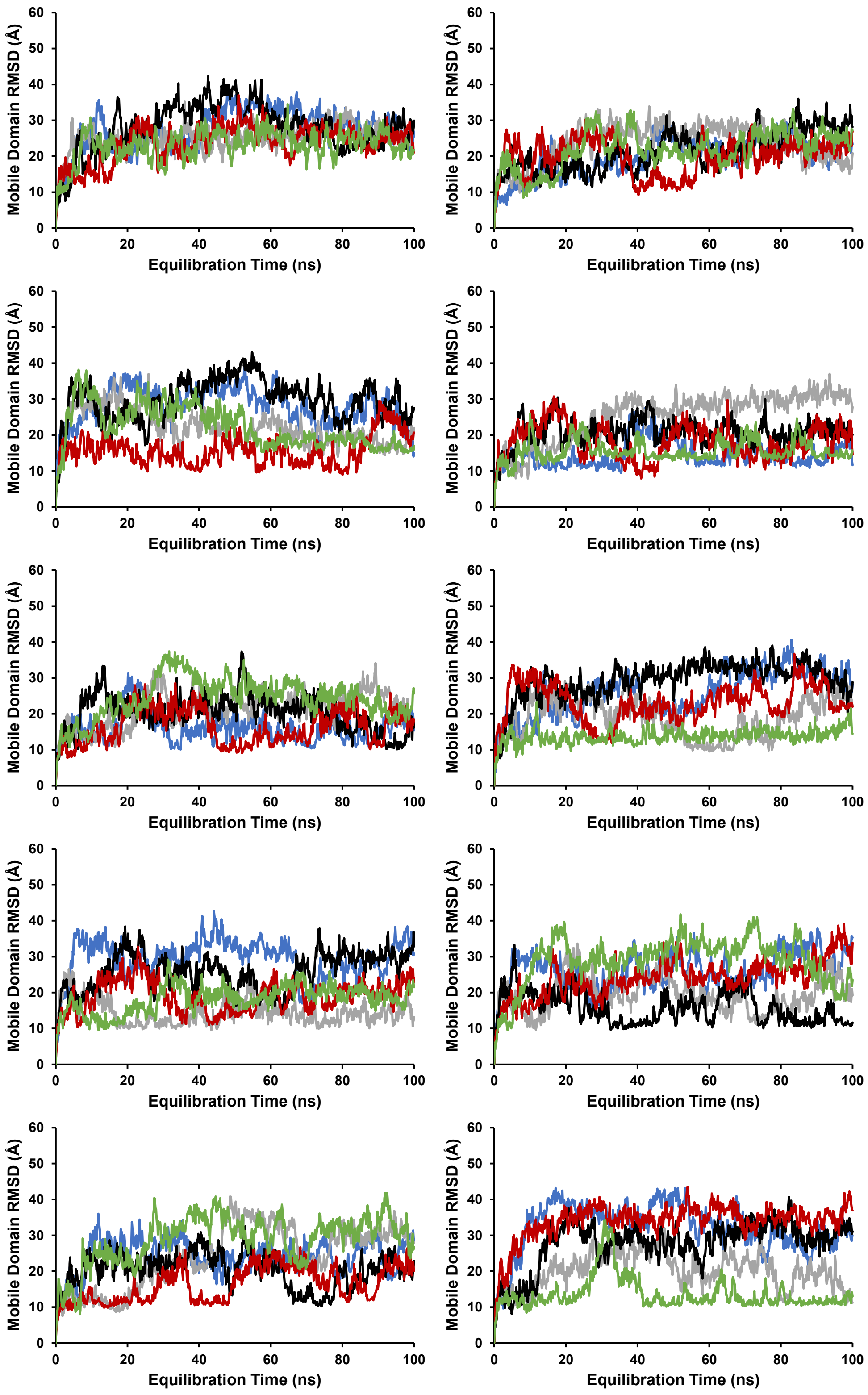


**Figure S8.** Traces of bound-state mobile domain RMSD vs equilibration time during MARTINI simulations for each replicate. Each graph shows equilibration trajectories for 5 replicates, each in a different color, for a total of 50 unique replicates across all 10 graphs.

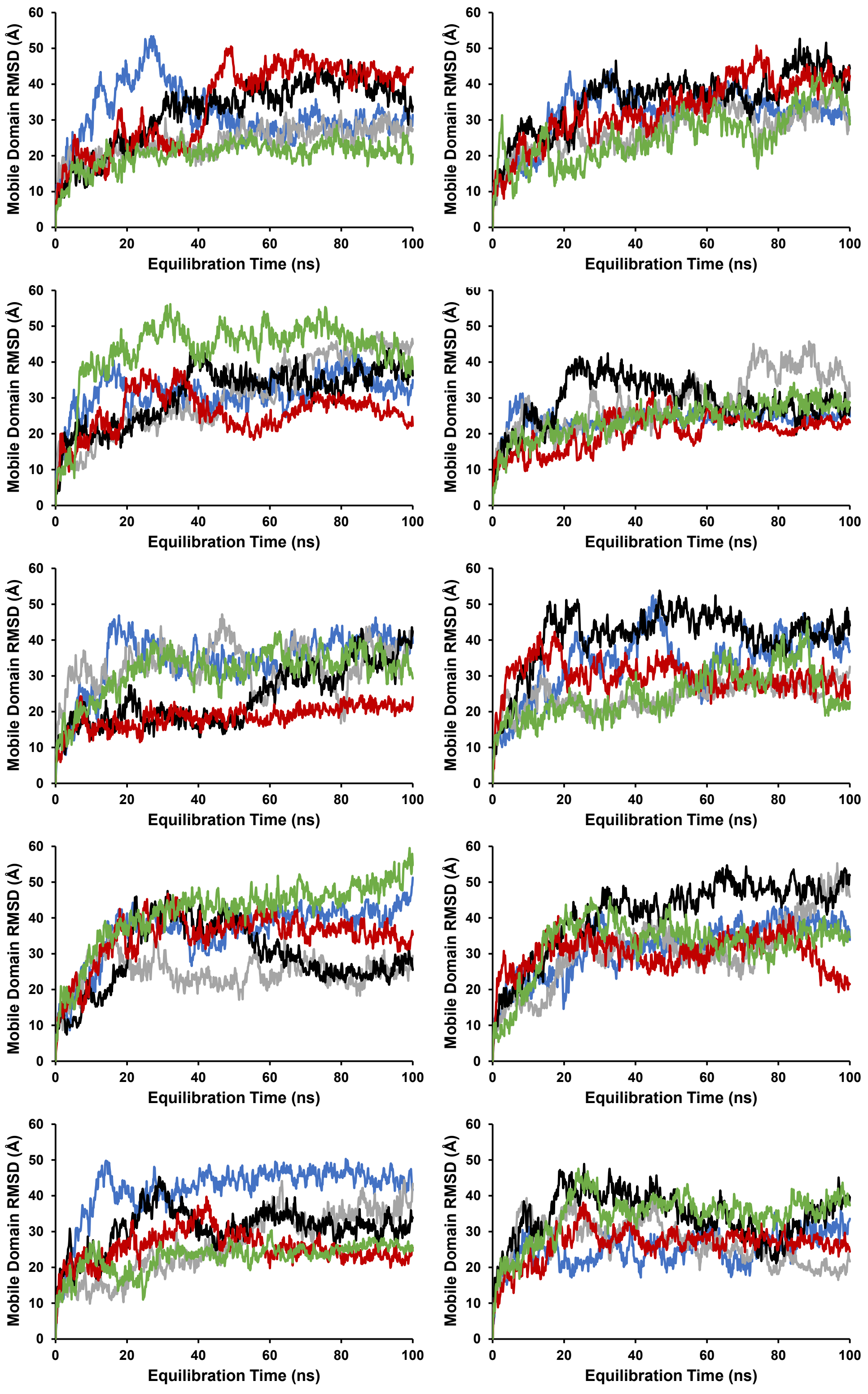


**Figure S9.** Traces of unbound-state mobile domain RMSD vs equilibration time during MARTINI simulations for each replicate. Each graph shows equilibration trajectories for 5 replicates, each in a different color, for a total of 50 unique replicates across all 10 graphs.

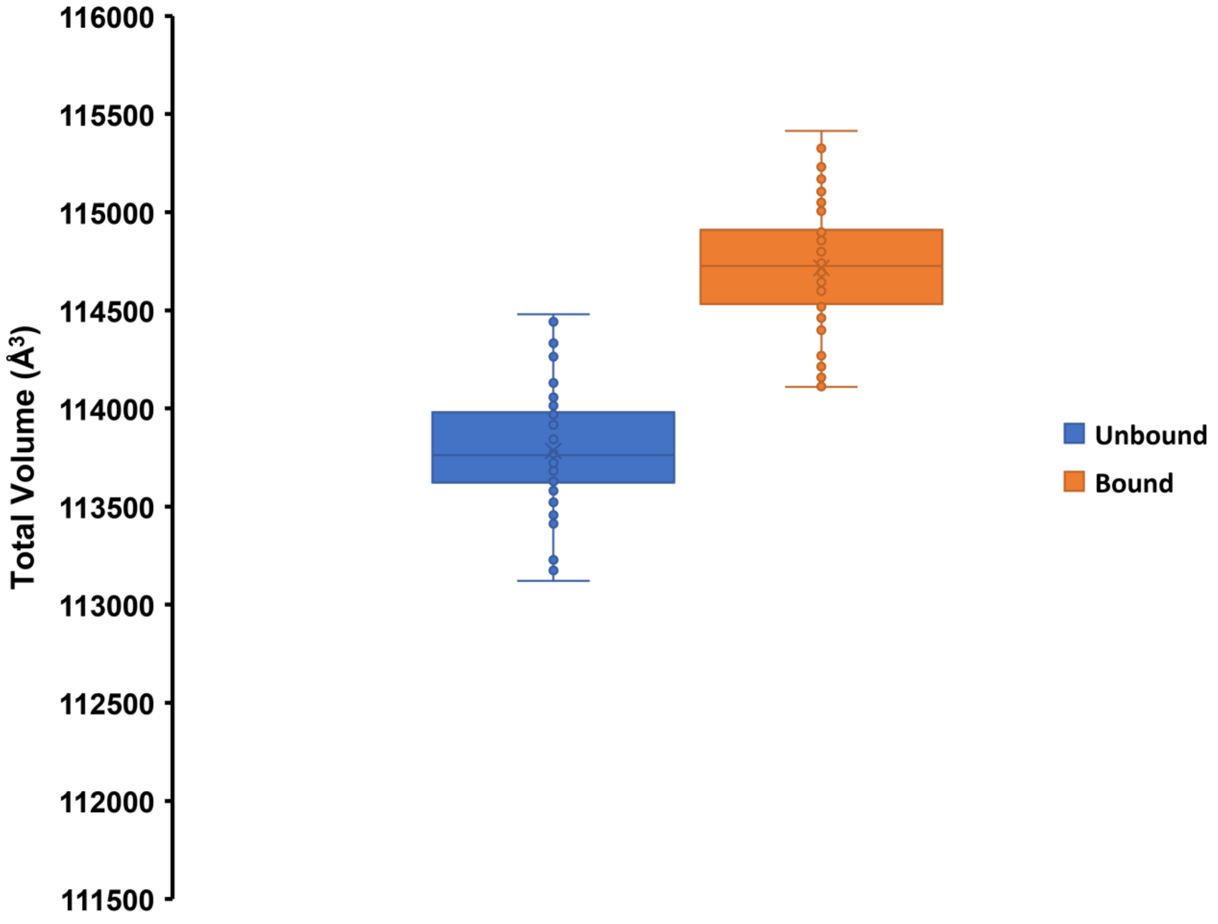


**Figure S10.** Box plots of solvent exclusion volume for unbound and bound states of the Cys-LeuFS conformational ensemble (n = 50 for each state). Volumes were calculated using ProteinVolume 1.3[32] and include both the van der Waals volume and internal cavity volume.

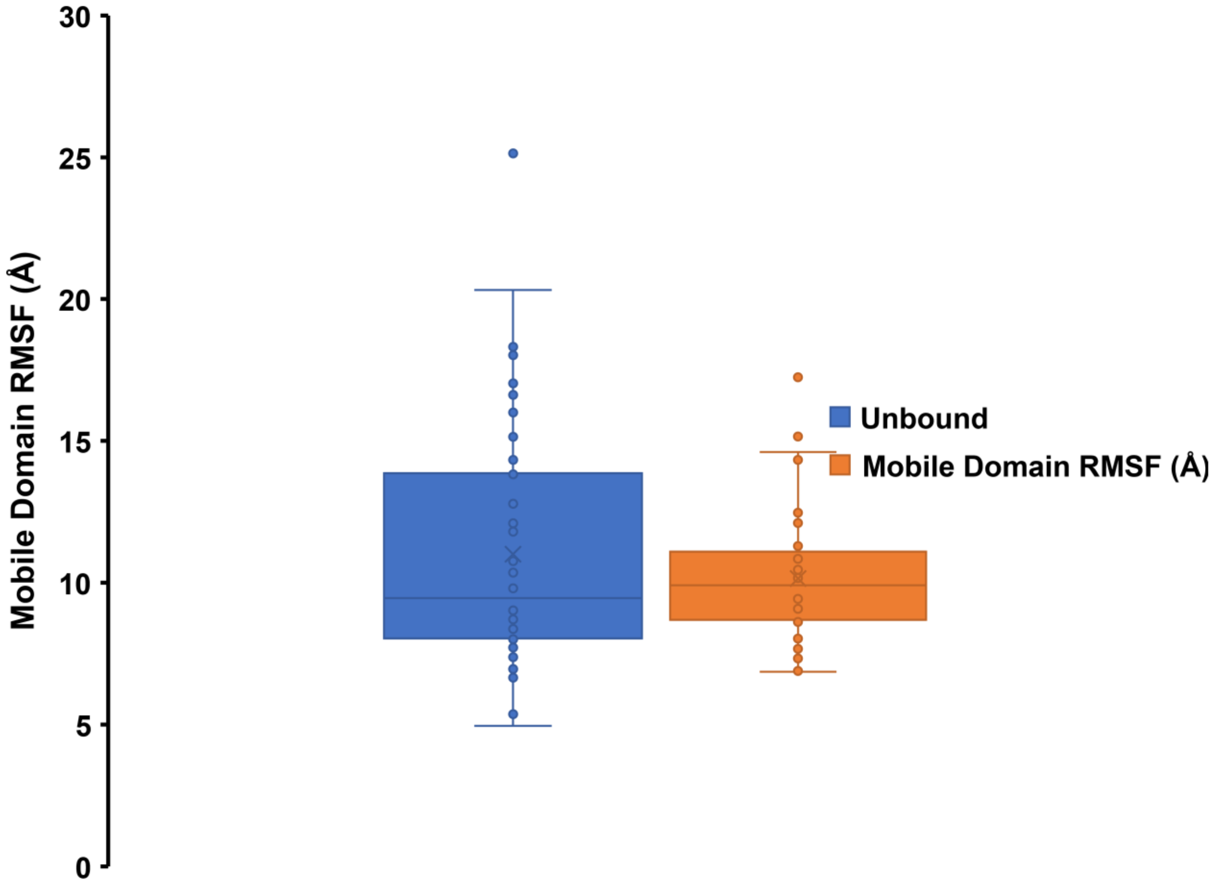


**Figure S11.** Box plots of mobile domain root mean square fluctuation (RMSF) for unbound and bound states of the Cys-LeuFS conformational ensemble (n = 50 for each state). Backbone (MARTINI BB bead) RMSF values were calculated using the final 50 ns of each simulation replicate and averaged over the mobile domain, following frame alignment using the static domain.

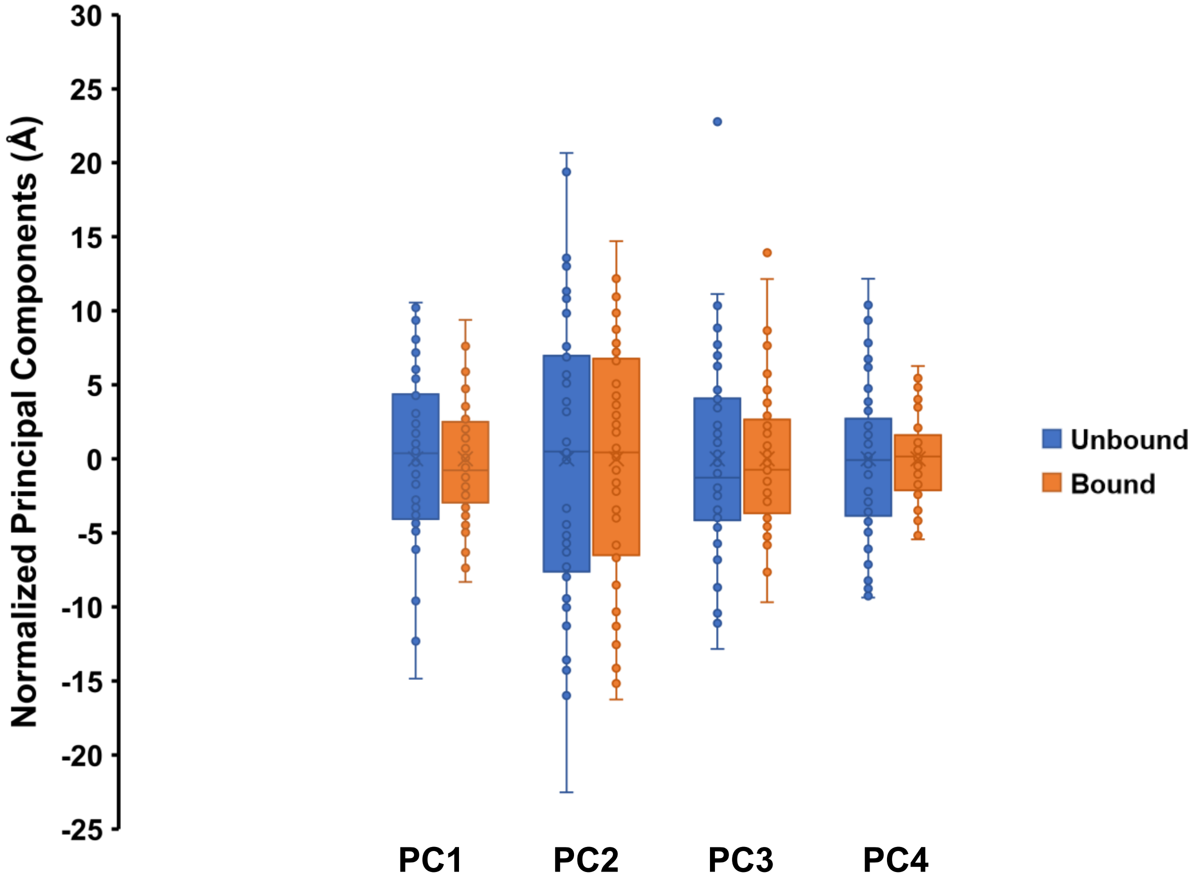


**Figure S12.** Box plots of average-normalized principal component projections for unbound and bound states of the Cys-LeuFS conformational ensemble (n = 50 for each state). The principal component analysis was carried out on the full 100-member ensemble. Following clustering of each principal component into unbound and bound state clusters, each cluster was normalized to an average value of 0 Å for comparison of distribution broadness.

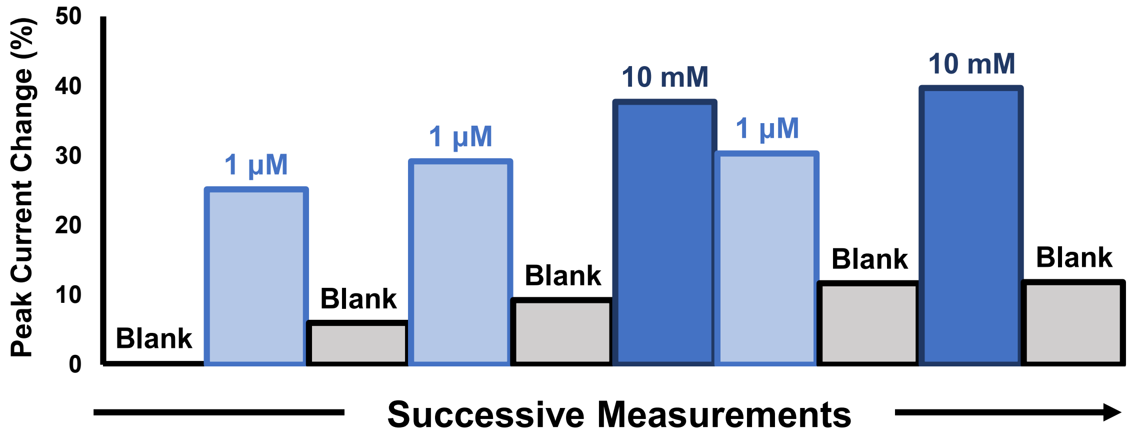


**Figure S13.** Reversibility of analyte binding by Cys-GlyFS was verified by successively cycling an Au screen-printed electrode (Au SPE) functionalized with Cys-GlyFS between solutions containing no (blank), 1 nM, and 10 mM glycine. Signal change was determined at the oxidation peak, relative to the initial blank.

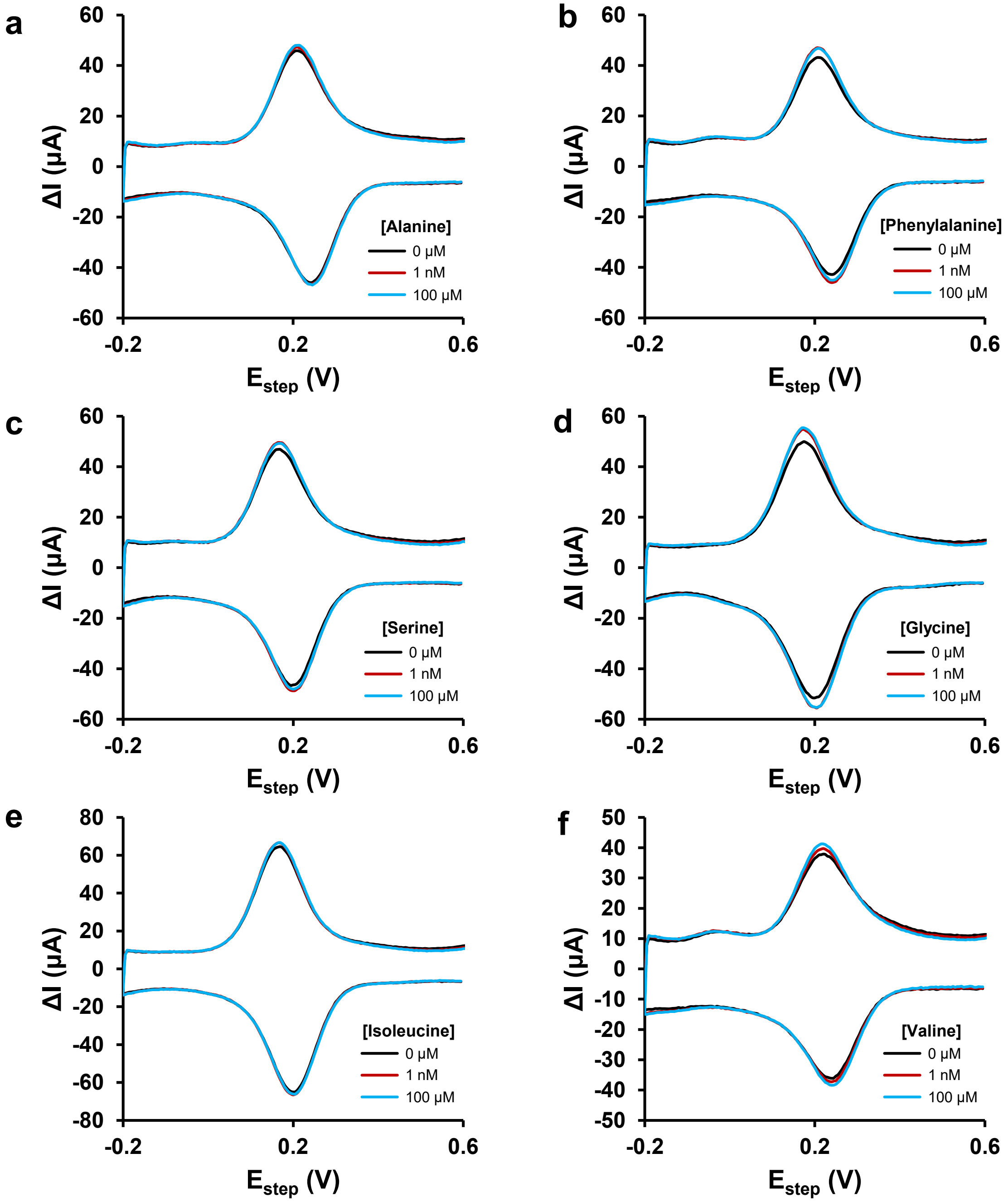


**Figure S14.** Cys-LeuFS selectivity towards leucine was validated through differential pulse voltammetry using was validated through differential pulse voltammetry using Au screen-printed electrodes (Au SPE) electrodes functionalized with Cys-GlyFS in solutions of 5 mM potassium ferricyanide containing one of six non-specific amino acid targets (alanine, phenylalanine, serine, glycine, isoleucine, and valine). Results are summarized in Figure 4.

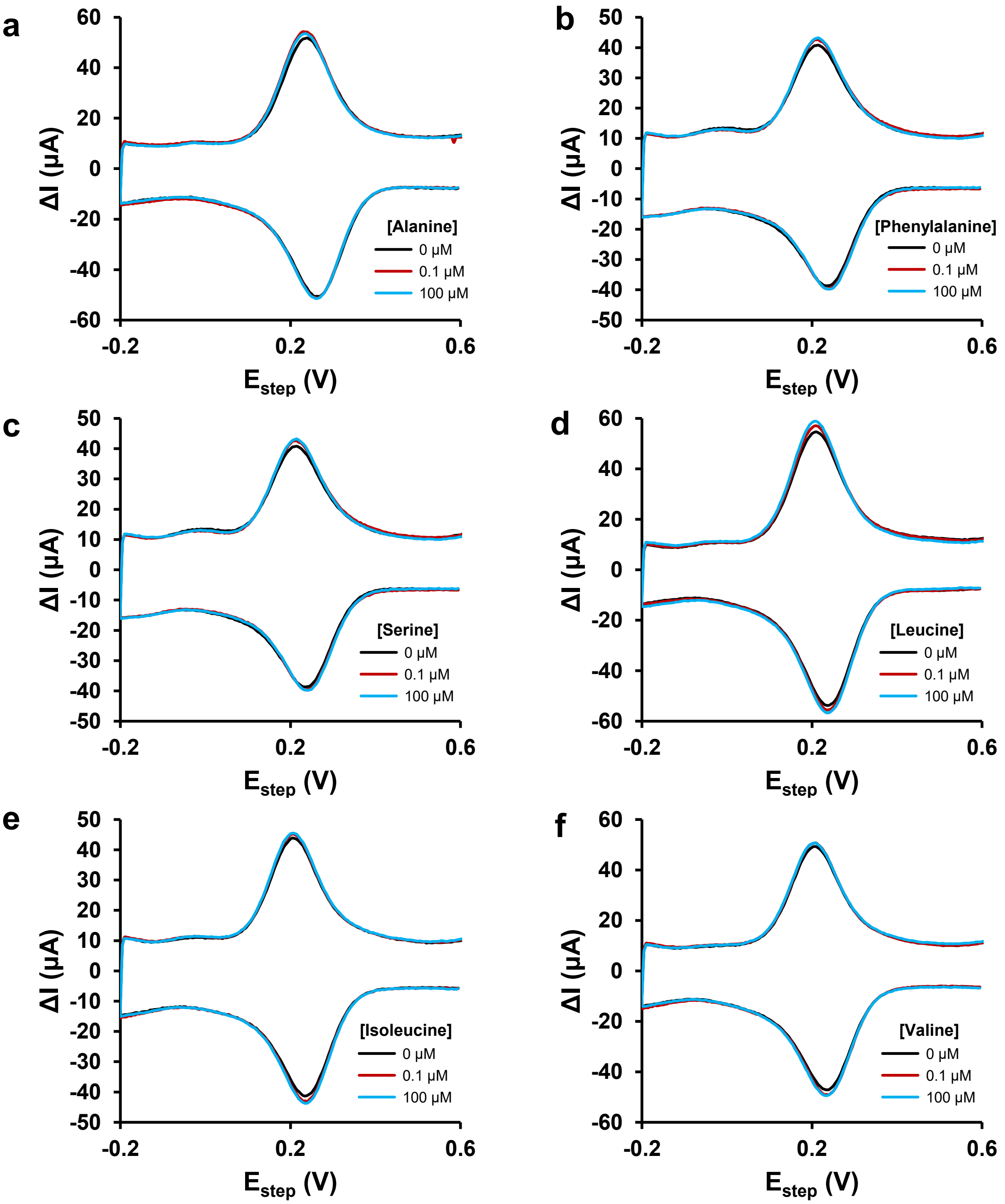


**Figure S15.** Cys-GlyFS selectivity towards glycine was validated through differential pulse voltammetry using Au screen-printed electrodes (Au SPE) electrodes functionalized with Cys-GlyFS in solutions of 5 mM potassium ferricyanide containing one of six non-specific amino acid targets (alanine, phenylalanine, serine, leucine, isoleucine, and valine). Results are summarized in Figure 4.
